# Searching Algorithm for Type IV Effector proteins (S4TE) 2.0: improved tools for type IV effector prediction, analysis and comparison

**DOI:** 10.1101/301150

**Authors:** Christophe Noroy, Thierry Lefrançois, Damien F. Meyer

## Abstract

Bacterial pathogens have evolved numerous strategies to corrupt, hijack or mimic cellular processes in order to survive and proliferate. Among those strategies, Type IV effectors (T4Es) are proteins secreted by pathogenic bacteria to manipulate host cell processes during infection. They are delivered into eukaryotic cells in an ATP-dependent manner via the type IV secretion system, a specialized multiprotein complex. T4Es contain a wide spectrum of features including eukaryotic-like domains, localization signals or a C-terminal translocation signal. A combination of these features enables prediction of T4Es in a given bacterial genome. In this study, we developed a web-based comprehensive suite of tools with a user-friendly graphical interface. This version 2.0 of S4TE (Searching Algorithm for Type IV Effector Proteins; http://sate.cirad.fr) enables accurate prediction and comparison of T4Es. Search parameters and threshold can be customized by the user to work with any genome sequence, whether publicly available or not. Applications range from characterizing effector features and identifying potential T4Es to analyzing the effectors based on the genome G+C composition and local gene density. S4TE 2.0 allows the comparison of putative T4E repertoires of up to four bacterial strains at the same time. The software identifies T4E orthologs among strains and provides a Venn diagram and lists of genes for each intersection. New interactive features offer the best visualization of the location of candidate T4Es and hyperlinks to NCBI and Pfam databases. S4TE 2.0 is designed to evolve rapidly with the publication of new experimentally validated T4Es, which will reinforce the predictive power of the algorithm. The computational methodology can be used to identify a wide spectrum of candidate bacterial effectors that lack sequence conservation but have similar amino acid characteristics. This approach will provide very valuable information about bacterial host-specificity and virulence factors, and help identify host targets for the development of new anti-bacterial molecules.

## INTRODUCTION

Proteobacteria have evolved specific effector proteins to manipulate host cell gene expression and processes, hijack immune responses and exploit host cell machinery during infection. These proteins are secreted by ATP-dependent protein complexes named type IV secretion systems (T4SS). Some T4Es have been identified and shown to be crucial for pathogenicity. To facilitate the identification of putative T4Es, we previously developed a bioinformatics tool called S4TE 1.0 (Searching Algorithm for Type IV secretion system effector proteins) [1].

In the present article, we present the second version of ‘S4TE’. S4TE 2.0 is a tool for *in silico* screening of proteobacteria genomes and T4E prediction based on the combined use of 14 distinctive features. In this updated version, modules searching for promoter motifs, homology, NLS, MLS and E-block are more efficient. A new module has been added in the workflow to locate phosphorylation (EPIYA-like) domains.

S4TE 2.0 consists of the S4TE 1.4 tool and a web interface available to non-commercial users at http://sate.cirad.fr. The web interface is designed to make S4TE 2.0 easy to use for biologists and more time efficient. Most of the genomes and plasmids available in the NCBI database of pathogenic bacteria that have a type IV secretion system have been loaded into the S4TE 2.0 database so effectors can be predicted in only a few clicks.

S4TE 2.0 offers advanced users an expert mode (S4TE-EM) they can use to customize S4TE 2.0 search parameters (*e.g.* exclude modules, modify module weightings). In this mode, S4TE 2.0 can be used as 14 independent programs to search for particular features in a given bacterial genome (*e.g.* NLS, C-ter charges).

A new function for comparative genomics (S4TE-CG) has been added to compare up to four predicted effectomes in just a few seconds.

All S4TE 2.0 results are interactive and linked to NCBI and Pfam databases.

## SOFTWARE AND ALGORITHM

### Programming

S4TE 2.0 software consists in a graphical interface (website) to use the S4TE 1.4 algorithm for genome analysis, Type IV effectors (T4Es) prediction and comparison of effectomes. S4TE 1.4 is an update of S4TE 1.0[1]. It is written in Perl programming language and uses NCBI, Pfam, EMBOSS, BioPerl and MitoFates libraries and its own proper programs and database. It was developed to improve the prediction performances of S4TE 1.0 and to provide new functionalities to search for new features, enable interactivity and comparative genomics. The 10 S4TE search modules in S4TE 1.0 were kept in S4TE 1.4. However, some modules have been modified (promoter motif search, homology, MLS, NLS, E-block and Pfam database) to improve their predictive power. A supplementary module (EPIYA search) has been added to the workflow. In this paper, only the EPIYA search module and the revised modules are described.

### Promoter motif search

As several T4Es in a given bacterium can be subjected to coordinated regulation with the same protein, *e.g.* PmrA[2], we used S4TE 2.0 to conduct a search for conserved motifs (potential regulatory motifs) in the short promoter regions of the genes. The aim was to improve S4TE 2.0 prediction of possible regulons of T4Es. Enriched DNA motifs were searched in a window of 100 nucleotides (nt) placed upstream of the start codon, using MEME[3]. Eight consensus motifs were identified in different bacteria (table 1). The corresponding motif search module of S4TE 2.0 extracts the 5’ Flanking intergenic regions (5’ FIRs) and searches for all these motifs thanks to a position-specific scoring matrix generated from multiple sequence alignments with the promoters of known T4Es. Only alignments with a score above the chosen threshold are selected. The threshold that yielded the highest sensitivity and specificity for each motif in the corresponding bacterium was chosen (Table 1).

**Table 1.** Enriched DNA motifs found in several bacteria in the 100 nucleotides upstream of known type IV effectors and implemented in S4TE 2.0 searches

| Name | Organism | Length | Threshold | Effector <sup>1</sup> | Non-effector <sup>2</sup> | Logo <sup>3</sup> |
| --- | --- | --- | --- | --- | --- | --- |
| PmrA | *Legionella* | 20 | 0.748 | 18.6 | 4.1 |  |
| Cpm | *Coxiella* | 20 | 0.87 | 18.2 | 0.02 |  |
| Cpm2 | *Coxiella* | 7 | 0.875 | 13.8 | 3.9 |  |
| Apm | *Anaplasma* | 14 | 0.7 | 77.8 | 9.9 |  |
| Apm2 | *Anaplasma* | 15 | 0.86 | 66.7 | 0.41 |  |
| Bapm | *Bartonella* | 19 | 0.75 | 62.5 | 2.2 |  |
| Hpm | *Helicobacter* | 20 | 0.68 | 1 | 0.39 |  |
| Bopm | *Bordetella* | 20 | 0.8 | 52.6 | 0.51 |  |

<sup>1</sup>Frequency of motif in effector promoters
<sup>2</sup>Frequency of motif in non-effector promoters
<sup>3</sup>Logo and motif were established using MEME software (Bailey TL *et al.*, 2009).

### Homology

BLAST 2.2 was used to compare proteins to search for homologies with known T4Es [4]. The cut-off of the S4TE 1.0 homology module was changed. S4TE 2.0 compares the database containing all known T4Es with the query proteome and returns all homologs with a cut-off of the expected value (E) <10-4. This E-value cut-off was selected to find real homologs between phylogenetically distant bacterial species. Databases containing proven effectors have also been updated (Table S1).

### Nuclear localization signals (NLS)

NLS are protein sequences that target proteins in the nucleus of eukaryotic cells[5]. We assume that the occurrence of NLS in a bacterial protein sequence would be a good indicator of secretion. There are two classes of NLS, monopartite and Bipartite. In S4TE 2.0, the search for monopartite NLS has been improved according to Ruhanen *et al.* [6]. We rewrote this module to add more known NLS motifs in the search. Monopartite NLS consist of [KR]-[KR]-[KR-][KR]-[KR], X-K-[KR]-[KRP]-[KR]-X, X-R-K-[KRP]-[KR]-X, X-R-K-X-[KR]-[KRP], X-K-[KR]-[KR]-X-[KRP], X-R-K-[KR]-X-[KRP], X-K-[KR]-X-[KR]-X-X, X-R-K-X-[KR]-X-X, X-K-[KR]-[KR]-X-X-X and X-R-K-[KR]-X-X-X motifs. Bipartite NLS were also searched with S4TE 1.0 motif (K-[KR]-X(6,20)-[KR]-[KR]-X-[KR]). The new module was tested with a dataset of 32 NLS and 32 no-NLS containing proteins (dataset 1). The module selected 24 true positives (TP) and only three false positives (FP). This represents a sensitivity (Se) of 75% and a specificity (Sp) of 91%.

### Mitochondrial Localization Signals (MLS)

MLS are signal sequences located in the N-terminus of proteins that are targeted to mitochondria. This sequence is cleaved after translocation of the protein inside the mitochondria[5,7]. To predict MLS in S4TE 2.0, we used the MitoFates tool[8]. MitoFates predicts mitochondrial presequences, a cleavable localization signal located in the N-terminal, and its cleaved position.

### E-block

The E-block domain consists of a glutamate sequence rich in C-terminal 30 amino acids and is associated with T4Es translocation in *L. pneumophila.* Huang *et al.* showed that an E-block motif is also important for the translocation of T4SS substrates[9]. In S4TE 2.0, the E-block module was modified according to Lifshitz *et al.* [10]. The E-block was searched in a window of 22 amino acids between position −4 C-terminal and −26 C-terminal. The motif that is searched for is a motif of 10 amino acids containing three or more glutamate (E) residues. The module was tested on 98 E-block and 98 no-E-block containing proteins (dataset 2). This module selected 60 TP and only 6 FP (Sensitivity of 61%, Specificity of 94%).

### Pfam database

The local Pfam database has been updated to find more eukaryotic domains of known effectors of *Legionella pneumophila*[10]. Eukaryotic domains were extracted from the whole Pfam database and added to the S4TE 2.0 workflow. All eukaryotic domains used for this search are listed in Table S2.

### EPIYA search

EPIYA search is a new module implemented in S4TE 2.0. The EPIYA domain is an eukaryotic phosphorylation motif[11]. In *H. pylori*, EPIYA has been shown to contribute to the secretion of a CagA effector[12]. We searched for conserved EPIYA motifs (EPIYA, ENIYE, NPLYE, EHLYA, TPLYA, EPLYA, ESIYE, EDLYA, EPIYG, EPVYA, VPNYA, EHIYD) in different bacteria that have a type IV secretion system and we searched for hypothetical EPIYA motifs using the motif E-X-X-Y-X.

### Validation

S4TE 2.0 is a software program with 14 independent modules. We tested all the modules independently. The 14 modules were weighted to make S4TE 2.0 efficient. The weighting of each module was calculated according to its performance in finding effectors in *L. pneumophila* Philadelphia I which has been shown to have the most extensive repertoire of T4Es ever identified, with 286 confirmed effectors [10].

Each module has its own weighting in S4TE 2.0 searches. The weightings were calculated for each module based on their Positive Predictive Value (PPV [PPV=TP/(TP+FP)]) for *L. pneumophila* (Table 2).

**Table 2.** Calculation of S4TE 2.0 weighting according to *Legionella pneumophila* Philadelphia 1 Positive Predictive Values (PPV) of each module

| S4TE 2.0 Features | 1 | 2 | 3 | 4 | 5 | 6 | 7 | 8 | 9 | 10 | 11 | 12 | 13 | 14 |
| --- | --- | --- | --- | --- | --- | --- | --- | --- | --- | --- | --- | --- | --- | --- |
| True Positives | 108 | 285 | 13 | 2 | 30 | 105 | 6 | 1 | 100 | 262 | 62 | 41 | 114 | 98 |
| False Positives | 434 | 34 | 27 | 106 | 101 | 783 | 79 | 6 | 231 | 2376 | 863 | 339 | 156 | 232 |
| PPV(%) | 20 | 89 | 32 | 2 | 23 | 12 | 7 | 14 | 30 | 10 | 7 | 10 | 42 | 30 |

The S4TE 2.0 prediction threshold was then defined to enable the best prediction by disregarding homology with known effectors. The threshold was chosen by examining the Sensitivity (Se), Specificity (Sp), Positive Predictive Value (PPV), Negative Predictive Value (NPV) and Accuracy (Acc) for thresholds ranging from 40 to 120 on the test dataset (Figure 2). The threshold was set at a score of 72 to obtain the global PPV possible with the least possible impact on sensitivity.

**Figure 1.**
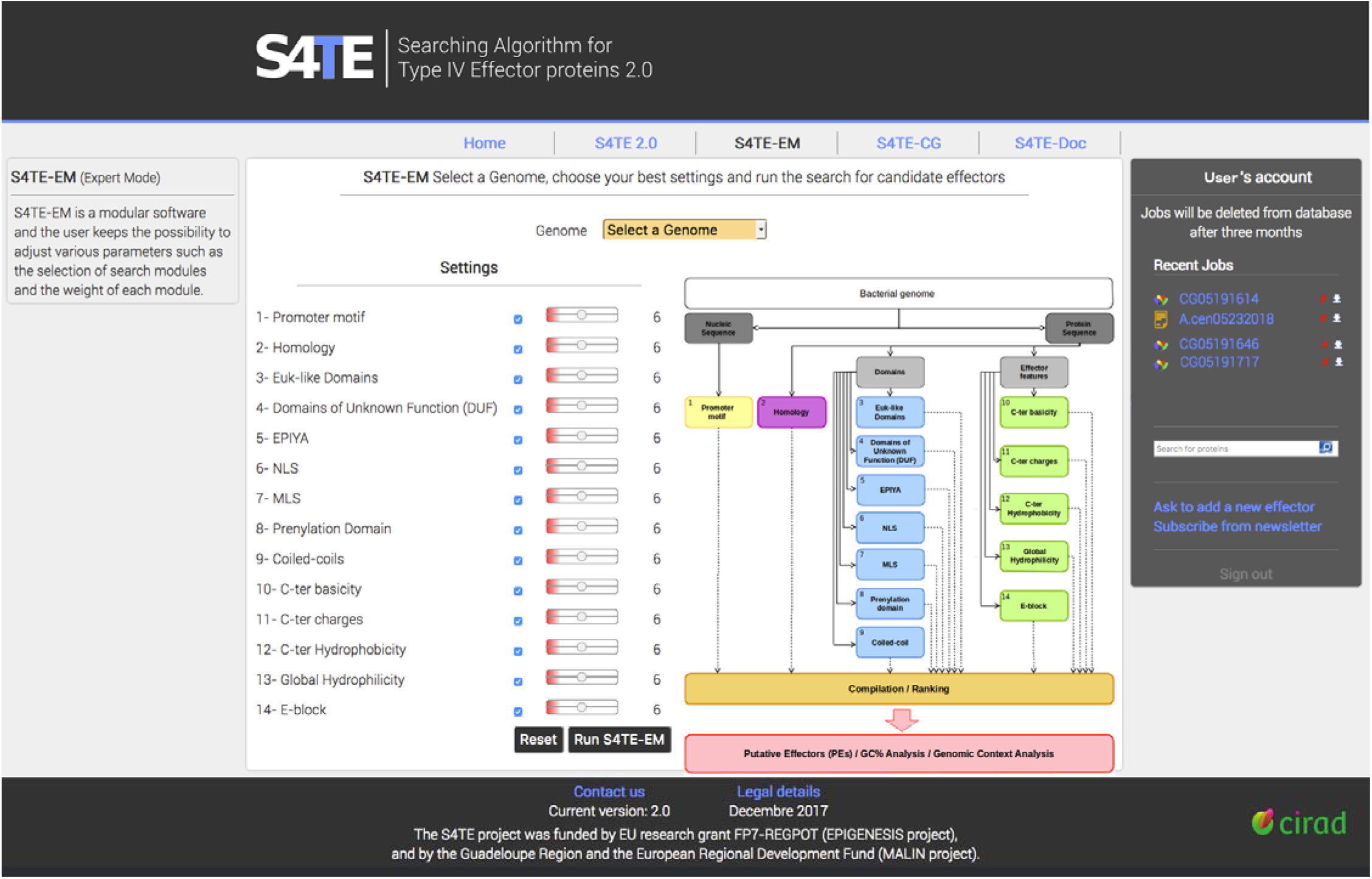
The new front page of the S4TE-EM tool. The right side provides some information about the page. The right side matches the user account. The user account shows all the jobs previously ran in S4TE 2.0 and S4TE-CG. This account makes it possible to search a protein with the search bar and to ask to add a proven T4 effector in the database. In the central part of the work space, the user can select a genome in the drop-down menu. In S4TE-EM, the user can change the weighting or disable one or more modules (on the left) shown in the S4TE diagram (on the right), and run S4TE-EM by clicking on the ‘Run S4TE-EM’ button.

**Figure 2.**
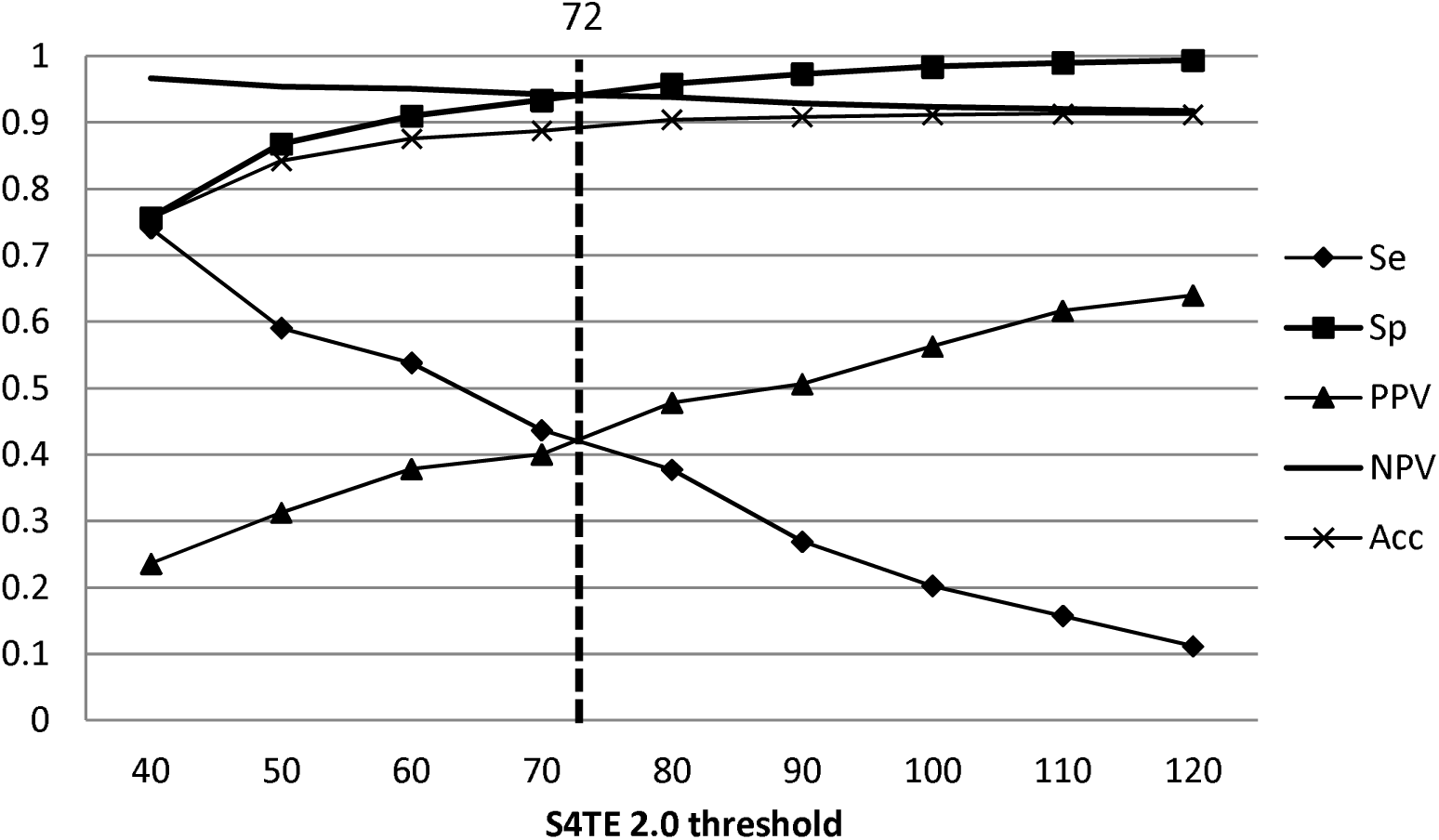
Distribution of S4TE 2.0 performances according to the threshold. Plot of the sensitivity (Se), specificity (Sp), positive predictive value (PPV), negative predictive value (NPV) and accuracy (Acc) of S4TE 2.0 with no homology module on *L. pneumophila* genome as a function of the S4TE 2.0 threshold. A threshold of 72 proved to be the best combination of these characteristics.

This threshold combined with weightings led to the correct prediction (true positives) of 282 of the 286 effectors of *L. pneumophila* (Se=98%, PPV=60%) and 96 incorrect predictions (false positives) (Sp = 96%, NPV = 99%).

With this update, S4TE 2.0 prediction is more powerful than that of S4TE 1.0 whose sensitivity was 14% lower. Without homology, sensitivity increased by 25% (data not shown). Other characteristics including specificity, accuracy and negative predictive value did not change significantly (table 3). S4TE 2.0 allows flexible, highly sensitive and specific detection of new putative T4SS effectors.

**Table 3.** Comparison between S4TE 1.0 and S4TE 2.0

| Software | S4TE1.0 |  | S4TE2.0 |  |
| --- | --- | --- | --- | --- |
| Homology | With | Without | With | Without |
| Sensitivity | 0.86 | 0.16 | 1 | 0.41 |
| Specificity | 0.97 | 0.97 | 0.93 | 0.93 |
| Positive Predictive Value | 0.74 | 0.44 | 0.60 | 0.43 |
| Negative Predictive Value | 0.98 | 0.91 | 1 | 0.94 |

### SATE-CG

S4TE-CG is a new tool designed to compare different repertoires of putative T4Es identified by S4TE 2.0. The corresponding S4TE-CG algorithm is described in Figure 3. The user can compare up to four effectomes simultaneously. S4TE 2.0 results from selected genomes (effectomes) are compared with Blastp 2.2 with an expected value (E) cut-off of <10-4 to find homologous proteins in each effectome. S4TE-CG successively compares all effectomes in a pairwise manner, the overlaps between the effectomes of each genome are calculated and the final results are plotted on a Venn diagram and listed in an interactive table. All effectors are clickable and the user is redirected to the S4TE 2.0 results on the effector concerned. The table can be easily copied and pasted for export.

**Figure 3.**
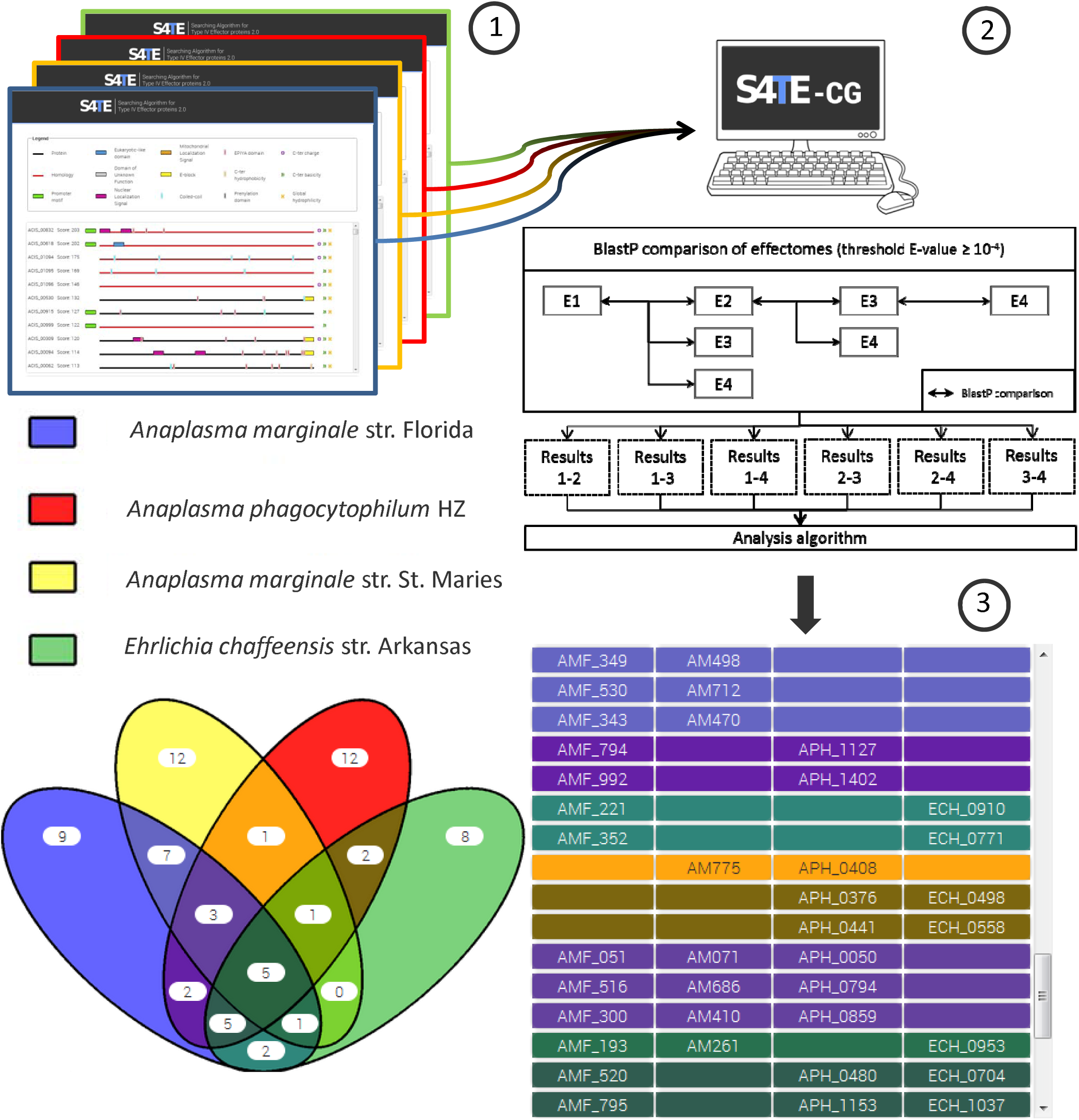
Flow chart of the comparison of 4 effectomes using S4TE-CG. Users can compare up to four genomes simultaneously. **1**. S4TE 2.0 results from selected genomes (effectomes) are compared with Blastp 2.2 to find homologous proteins in each effectome. **2**. S4TE-CG successively compares all effectomes in a pairwise manner, and calculates any overlaps between the effectomes of each genome. **3**. The final results are plotted on a Venn diagram and listed in an interactive table.

### Software availability

S4TE 2.0 is a web interface and the S4TE 1.4 package is freely available to non-commercial users at http://sate.cirad.fr/S4TE-Doc.php. All programming was done using Perl 5.18 and BioPerl 1.6.1. The software runs on Linux platforms (Ubuntu 14.04 and Mac OS X). All required packages and the installation process are described in the user guide included in the package. The user guide also details S4TE options for running S4TE. By default, the command line to launch S4TE is `./S4TE.pl -f “Genbank_file”` in the S4TE folder (`cd way_to_S4TE/S4TE/`). Some options are available for the user to launch S4TE: `-c`, suppression of a module in the pipeline; `-w`, modification of the weight of each module in the pipeline; `-t`, imposition of a threshold for effector selection. Each S4TE module creates a `.txt` file in the folder `way_to_S4TE/S4TE/Jobs/job<Name_of_genome_folder><year><month><day><hour><min>` All the results are compiled in the *CompilationFile.txt* and *Results.txt* in the same folder.

## WEB INTERFACE

### Design and general features

The S4TE 2.0 website is powered from scratch on the ‘CIRAD web server’. All the features of the web site were tested on common web browsers. S4TE 2.0 found T4Es in large genome databases (Table S3) available to all users. A user account is available and necessary to keep your jobs up to three months, to import your own genome in a S4TE 2.0 temporary database and to ask to add a new proved effector in the database. The addition of an effector to the database must be accompanied by a reference (scientific article) and will be checked manually before the effector is added to the database. Those who subscribe to the newsletter will be notified by email about the addition of new effectors to the database and the effector will be visible in the S4TE 2.0 tab strip. This free account allows users to search for proteins in the S4TE 2.0 database using the name, the locus tag or NCBI number of a protein in the search bar. The account also allows the user to subscribe to the S4TE newsletter that summarizes any changes made to the software, and provide updates on the latest research on Type IV Effectors.

### S4TE 2.0 is a simple and user-friendly tool

S4TE 2.0 is a web-based user-friendly tool that gets results in only a few clicks. The user can locate a chromosome in more than 340 bacterial genomes and plasmids available in the database and the results can be viewed by clicking on run S4TE 2.0 (Table S3). If the desired genome is not available in the databases, the user can import it with a GenBank file (.gbk). S4TE 2.0 will import the file to a temporary database for three months. All S4TE tools (S4TE-EM and S4TE-CG) can then be used on the genome by the owner. The S4TE 2.0 web page allows users to read some of the news published in the newsletter. Five news items are visible on the S4TE2.0 web page, but all the news can be found by clicking on the bottom right link.

Figure 4 presents some results obtained with S4TE 2.0. All the proteins in the selected genome are represented on the S4TE 2.0 web results page. A score was calculated for each protein based on the weighting of each module. Proteins were ranked according to the same score. All proteins whose scores are above the threshold are considered as belonging to the S4TE 2.0 effectome. An iconography was created to help read the list (Figure 4A). Users can find all the details concerning each characteristic of a given protein by clicking on the protein concerned on the web results page.

**Figure 4.**
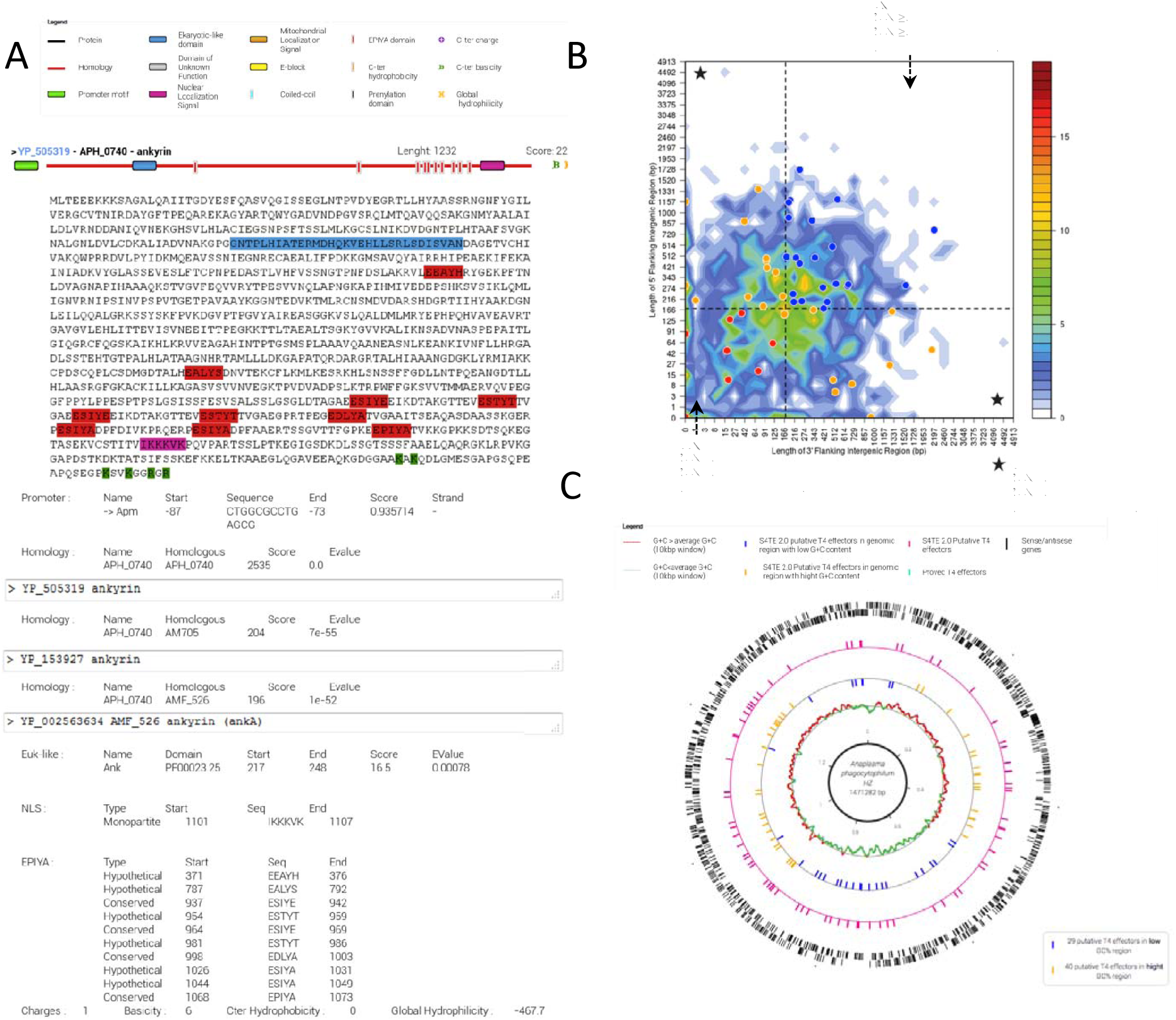
Example of S4TE 2.0 results for *Anaplasma phagocytophilum* HZ. APH-0740. **A**. Schematic representations of proteins with different characteristics present in the sequence are shown. Characteristics are easy to find by highlighting the corresponding sequence in the effector sequence. These characteristics are detailed below the sequence. **B**. Distribution of S4TE 2.0 predicted type IV effectors (T4Es) according to local gene density. The predicted T4Es are plotted according to the length of their flanking intergenic regions (FIRs). All *A. phagocytophilum* genes were sorted into 2-dimensional bins according to the length of their 5′ (y-axis) and 3′ (x-axis) FIRs. The number of genes in the bins is represented by a color-coded density graph. Genes whose FIRs are both longer than the median FIR length were considered as gene-sparse region (GSR) genes. Genes whose FIRs are both below the median value were considered as gene-dense region (GDR) genes. In-between region (IBR) genes are genes with a long 5′FIR and short 3′FIR, or inversely. Candidate T4Es predicted using the S4TE2.0 algorithm were s plotted on this distribution according to their own 3′ and 5′ FIRs. A color is assigned to each of the three following groups: Red to GDRs, orange to IBRs, and blue to GSRs. **C**. Genome-wide distribution of predicted effectome according to the G+C content. From outer track to inner track, sense and antisense genes (black), S4TE 2.0 putative T4Es (pink), proved T4Es (turquoise), S4TE 2.0 putative T4Es in genomic region with low G+C content (yellow), S4TE 2.0 putative T4Es in genomic region with high G+C content (blue), G+C e average G+C (red), G+C < average G+C (green).

When a user runs S4TE 2.0, in addition to the results page, two graphs are automatically drawn. The first shows the distribution of predicted effectors according to local gene density (Figure 4B). The second one displays the distribution of predicted T4Es according to the G+C content along the genome (Figure 4C).

### S4TE-EM Expert mode for accurate searching

S4TE-EM is the expert mode of S4TE 2.0. S4TE-EM allows the user to modify the weights of each module and to deactivate one or more modules in the search (Figure 1). The weight of a module can be changed by moving the position of the cursor next to the name of each module. Weightings can be changed between the lowest weighting available for the module and the threshold of S4TE 2.0 (`t`=72). The lowest weight is calculated independently for each module as a function of the positive predictive value and corresponds to a value equal to 0.5. Users can also cancel one or more modules in the pipeline by unchecking the box next to the name of the module (Figure 1).

All the modules are independent and users can use S4TE-EM to locate the same characteristic throughout the genome. For example, if the user disables all the modules except NLS, S4TE-EM will find all proteins with an NLS in the genome, meaning users can use S4TE-EM as a new genome analysis tool.

### S4TE-CG Comparative genomics to compare effectomes

S4TE-CG is a new tool designed to compare different effectomes predicted by S4TE 2.0. Users can choose up to four effectomes in S4TE 2.0 databases or upload a genome present in the temporary database. S4TE-CG displays results in a Venn diagram and in an interactive table. Users can easily find different subsets of information in the appropriate table by referring to the different colors in the Venn diagram (Figure 3). Information about each effector can easily be found by clicking on the name of the effector in the table. Or users can simply copy and paste the table in a .csv file.

## CONCLUSION

This paper presents updated S4TE software. The computational tool is designed to predict the presence of T4SS effector proteins in bacteria. The identification of T4Es and some characteristics are improved in this update. Compared with a machine learning approach, using S4TE 2.0 to predict T4Es in *Legionella* and *Coxiella* species[10,13,14] improved sensitivity (98% for S4TE 2.0 and 89% for Wang *et al.)* and equivalent specificity (97% for Wang *et al.* and 93% for S4TE 2.0). S4TE 2.0 is easy to use. Only an internet connection and a few clicks are needed to search for T4Es in more than 340 bacterial genomes and plasmids. The results are displayed instantaneously for easy reading. An automated pipeline is also provided to analyze and visualize effector distribution in the genome according to G+C content and local gene density. S4TE 2.0 results are linked to bioinformatics databases like NCBI and Pfam. The S4TE 2.0 database is designed to evolve and will be updated by adding new proven effectors and new bacterial genomes. S4TE 2.0 not only predicts the T4Es but also their subcellular localization (NLS, MLS, prenylation) and the function of these proteins (Coiled coils, EPIYA, Euk-like, etc.). All these features make S4TE 2.0 a powerful software for studies of T4Es.

S4TE 2.0 also offers an expert mode, which allows users to make manual adjustments to the weight of the modules. Each module that searches for a feature or a characteristic can be used independently. S4TE EM can be viewed and use as 14 independent programs. This could facilitate the annotation of new genomes by looking for specific features such as NLS, prenylation domains, etc.

Finally, S4TE-CG makes it possible for users to compare effectomes to highlight core T4 effectomes and/or accessory T4 effectomes to understand how effectomes evolved, and may provide clues to the specificity of different strains.

